# Large-scale changes to mRNA polyadenylation in temporal lobe epilepsy

**DOI:** 10.1101/725325

**Authors:** Alberto Parras, Laura de Diego-Garcia, Mariana Alves, Edward Beamer, Giorgia Conte, James Morgan, Ivana Ollà, Yasmina Hernandez-Santana, Norman Delanty, Michael A. Farrell, Donncha F. O’Brien, David C. Henshall, Raúl Méndez, José J. Lucas, Tobias Engel

**Author notes:** Correspondence: Tobias Engel, Ph.D., Department of Physiology and Medical Physics, Royal College of Surgeons in Ireland, 123 St. Stephen’s Green, Dublin 2, Ireland, José J. Lucas, Centro de Biología Molecular ‘Severo Ochoa’ (CBMSO) CSIC/UAM, C/ Nicolás Cabrera, 1, Campus UAM de Cantoblanco, 28049 Madrid, Spain. These authors contributed equally (Alberto Parras, Laura de Diego-Garcia).

## Abstract

The molecular mechanisms that shape the gene expression landscape during the development and maintenance of chronic states of brain hyperexcitability are incompletely understood. Here we show that cytoplasmic mRNA polyadenylation, a posttranscriptional mechanism for regulating gene expression, undergoes widespread reorganisation in temporal lobe epilepsy. Specifically, over 25% of the hippocampal transcriptome displayed changes in their poly(A) tail in mouse models of epilepsy, particular evident in the chronic phase. The expression of cytoplasmic polyadenylation binding proteins (CPEB1-4) was found to be altered in the hippocampus in mouse models of epilepsy and temporal lobe epilepsy patients and CPEB4 target transcripts were over-represented among those showing poly(A) tail changes. Supporting an adaptive function, CPEB4-deficiency leads to an increase in seizure severity and neurodegeneration in mouse models of epilepsy. Together, these findings reveal an additional layer of gene expression control during epilepsy and point to novel targets for seizure control and disease-modification in epilepsy.

## Introduction

Epilepsy is one of the most common chronic neurological disorders, affecting approximately 70 million people worldwide^1,2^. Temporal lobe epilepsy (TLE) is the most common refractory form of epilepsy in adults and typically results from an earlier precipitating insult that causes structural and functional reorganisation of neuronal-glial networks within the hippocampus resulting in chronic hyperexcitability^3^. These network changes, which include selective neuronal loss, gliosis and synaptic remodelling, are driven in part by large-scale changes in gene expression^4-7^. The gene expression landscape continues to be dysregulated once epilepsy is established^8^.

Recent studies have uncovered important roles for post-transcriptional mechanisms during the development of epilepsy. These include the actions of small noncoding RNA, such as microRNA and post-translational control of protein turnover via the proteasome contributing to altered levels of ion channels, changes in neuronal micro- and macro-structure and glial responses within seizure-generating neuronal circuits^9-13^. The molecular mechanisms underlying the transcriptional and translational landscape in epilepsy remain, however, incompletely understood.

In the cell nucleus, the majority of mRNAs acquire a non-templated poly(A) tail. Although the addition of a poly(A) tail seems to occur by default, the subsequent control of poly(A) tail length is highly regulated both in the nucleus and cytoplasm^14^. Cytoplasmic mRNA polyadenylation contributes to the regulation of the stability, transport and translation of mature transcripts, and is therefore an essential post-transcriptional mechanism for regulating spatio-temporal gene expression^14^.

The cytoplasmic polyadenylation element binding proteins (CPEBs) are sequence-specific RNA-binding proteins (RNABPs) and key regulators of mRNA translation via the modulation of poly(A) tail length^15^. The CPEB family is composed of four members in vertebrates (CPEB1-4) with CPEB2-4 sharing more homology than with CPEB^16^. To function, CPEBs bind to cytoplasmic polyadenylation element (CPE) sequences, located at the 3’ untranslated region (3’UTR) of the target mRNAs. They nucleate a complex of factors that regulate poly(A) tail length, thereby controlling both translational repression and activation^15^. In the brain, CPEBs mediate numerous cellular processes including long-term potentiation, synaptic plasticity and expression of neurotransmitter receptors^17-19^. Functional roles for CPEBs in brain diseases include CPEB4 a critical regulator of risk genes associated with autism^21^ and as protective against ischemic insults *in vivo*^20^. Notably, evoked seizures result in increased CPEB4 expression^22^ and mice lacking Cpeb1 and the *Fmr1* gene display decreased susceptibility to acoustic stimulation-induced seizures^23^. Here, we explored changes in mRNA polyadenylation in experimental mouse models of epilepsy, revealing large-scale alterations as a major feature of the gene expression landscape in epilepsy.

## Results

### Genome-wide mRNA polyadenylation profiling reveals deadenylation of epilepsy-related genes

To investigate whether seizures and epilepsy impact on mRNA polyadenylation, we used the well characterized intraamygdala kainic acid (KA)-induced status epilepticus mouse model^24^. In this model of acquired epilepsy, an intraamygdala microinjection of KA leads to status epilepticus and wide-spread neurodegeneration involving the ipsilateral cortex and CA3 subfield of the hippocampus. All mice treated with intraamygdala KA develop epilepsy following a short latency period of 3-5 days, experiencing 2-5 seizures per day^24^.

We first used mRNA microarrays to map both, changes in the rate of gene transcription and potential genome-wide alterations in poly(A) tail length in the ipsilateral hippocampus at two time-points; 8 hours (h) post-status epilepticus (acute injury) and at 14 days post-status epilepticus (epilepsy) (Fig. 1a). Changes in poly(A) tail length were analyzed via poly(U) chromatography. Here, by using a differential elution with either 25% or 90% formamide, total hippocampal mRNA extracts were separated into two fractions, one enriched in mRNAs with a short and one enriched in mRNAs with a long poly(A) tail, respectively^21^ (Fig. 1b). This approach revealed that large-scale increases in mRNA levels accompany the early phase after status epilepticus (4991 genes) when compared to established epilepsy (968 genes) (Fig. 1b). Therefore, regulation of mRNA levels seems to dominate the altered gene expression landscape immediately following status epilepticus which is later reduced in epilepsy. In sharp contrast to alterations in transcript levels, changes in poly(A) tail length were much more pronounced in samples from mice with established epilepsy, affecting 28.6% of the total mRNA pool (6177 genes) when compared to status epilepticus affecting 9.6% of the genome (2088 genes) (Fig. 1b).

**Fig. 1.**
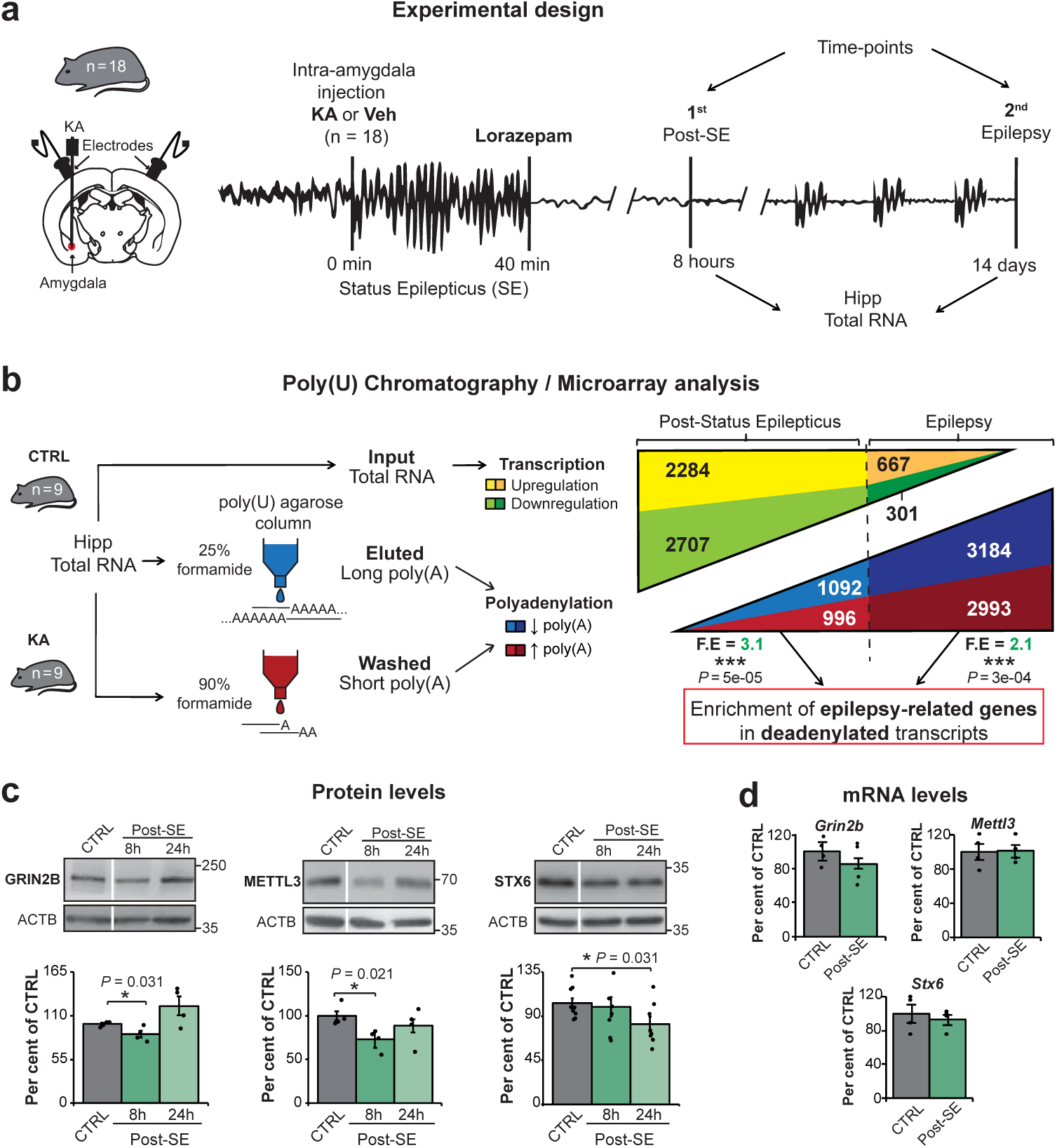
Genome-wide mRNA polyadenylation changes following status epilepticus and during epilepsy. **a,** Schematic showing the experimental design using the intraamygdala KA mouse model of acquired epilepsy. Hippocampi were collected at two time-points: 8 hours post-status epilepticus (acute injury, n = 9) and 14 days post-status epilepticus (epilepsy, n = 9). KA, kainic acid; SE, status epilepticus. **b,** Experimental design of poly(U) chromatography and microarray analysis (left) and comparison between genes with dysregulated transcription vs. mRNA polyadenylation changes (right) (Total amount of analyzed genes = 21566). F.E, fold enrichment. **c,** Protein levels in the ipsilateral hippocampus of WT mice injected with vehicle (Control) vs. KA (status epilepticus) at 8 h and 24 h post-status epilepticus of GRIN2B (n = 4), METTL3 (n = 4) and STX6 (n = 8). CTRL, control; WT, wildtype; SE, status epilepticus. **d,** mRNA levels in the ipsilateral hippocampus at 8 h post-status epilepticus, vehicle vs. KA-treated mice (n = 4). Data were analyzed and =normalized to the expression of ACTB **b,** One-sided Fisher’s exact test, *P* values of deadenylated epilepsy-realted genes versus total deadenylated transcripts. **c, d,** Two-sided unpaired t-test. Data are mean ± S.E.M. 95% CIs. \**P* < 0.05.

In order to identify functional groups of genes and pathways affected by changes in polyadenylation, transcripts with poly(A) tail alterations were analysed by Gene Ontology (GO) terms using the bioinformatic tool DAVID^25^. Notably, pathways related to epilepsy were particularly abundant among genes undergoing a shortening in their poly(A) tail following status epilepticus and during epilepsy (Extended Data Fig. 1a). Shortening of selected deadenylated transcripts was validated by high-resolution poly(A) tail (Hire-PAT) assay (Extended Data Fig. 1b). To explore whether changes in mRNA polyadenylation disproportionately affect genes implicated in the pathogenesis of epilepsy, we compiled a set of epilepsy-related genes from three recent independent studies. These studies included: a) genes where mutations cause epilepsy^26^, b) genes with ultra-rare deleterious variations in familial genetic generalized epilepsies (GGE) and non-acquired focal epilepsies (NAFE)^27^ and c) genes localized in loci associated with epilepsy^28^ (Supplementary Table 2). Remarkably, transcripts displaying poly(A) tail shortening showed an enrichment in genes implicated in epilepsy at both time-points, post status epilepticus and during epilepsy (Fig. 1b). This enrichment persisted after comparing our dataset with genes specifically expressed in the brain, thus proving this enrichment to be specific for epilepsy-related genes (Extended Data Fig. 1c). Genes identified in our array to undergo mRNA deadenylation following status epilepticus, also showed a reduction in their protein levels, demonstrating that mRNA deadenylation leads to reduced protein expression. This included the Glutamate ionotropic receptor NMDA type subunit 2B (GRIN2B), previously shown to play a role during epilepsy^26^, Syntaxin 6 (STX6) and N6-adenosine-methyltransferase (METTL3), genes not associated with epilepsy before (Fig. 1c). No significant changes in transcript levels of our selected genes was observed when analysed post-status epilepticus, further suggesting this decrease in expression is due to mRNA deadenylation (Fig. 1d). Previous *in vivo* studies showed that deadenylation, by resulting in decreased protein output, is more disruptive than poly(A) tail elongation in the brain^21^. Our results therefore indicate that deadenylation may play a role during the development of epilepsy through diminishing the translation of specific target genes.

Taken together, our results demonstrate changes in mRNA polyadenylation following seizures and epilepsy affecting over 25% of the total transcriptome with deadenylation disproportionately affecting transcripts encoding proteins related to epilepsy. Our results, therefore, reveal mRNA polyadenylation as a previously unrecognized layer of gene expression control in epilepsy.

### Increased expression of CPEBs in the hippocampus of TLE patients

To identify possible candidate RNA Binding Proteins (RNABPs) responsible for driving seizure-induced alterations in poly(A) tail length, we first compiled a list of genes whose transcripts are known to bind to the main RNABPs involved in cytoplasmic polyadenylation^29^ including human antigen R (HUR), which prevent deadenylation^30^; PUMILIO, which promotes deadenylation^31^ and CPEBs^21^ (Fig. 2a and Supplementary Table 3). This analysis revealed that CPEB binders were highly enriched among genes that showed poly(A) tail alterations following status epilepticus and during epilepsy (Fig. 2b). Interestingly, binders of CPEB1 and of CPEB4 (the latter representing the CPEB2-4 subfamily) have been identified in the brain^21^; while targets of both CPEBs were increased among genes with poly(A) tail changes, the percentage of CPEB4 binders was significantly higher when compared to CPEB1 binders, particularly during epilepsy (Extended Data Fig. 2a).

**Fig. 2.**
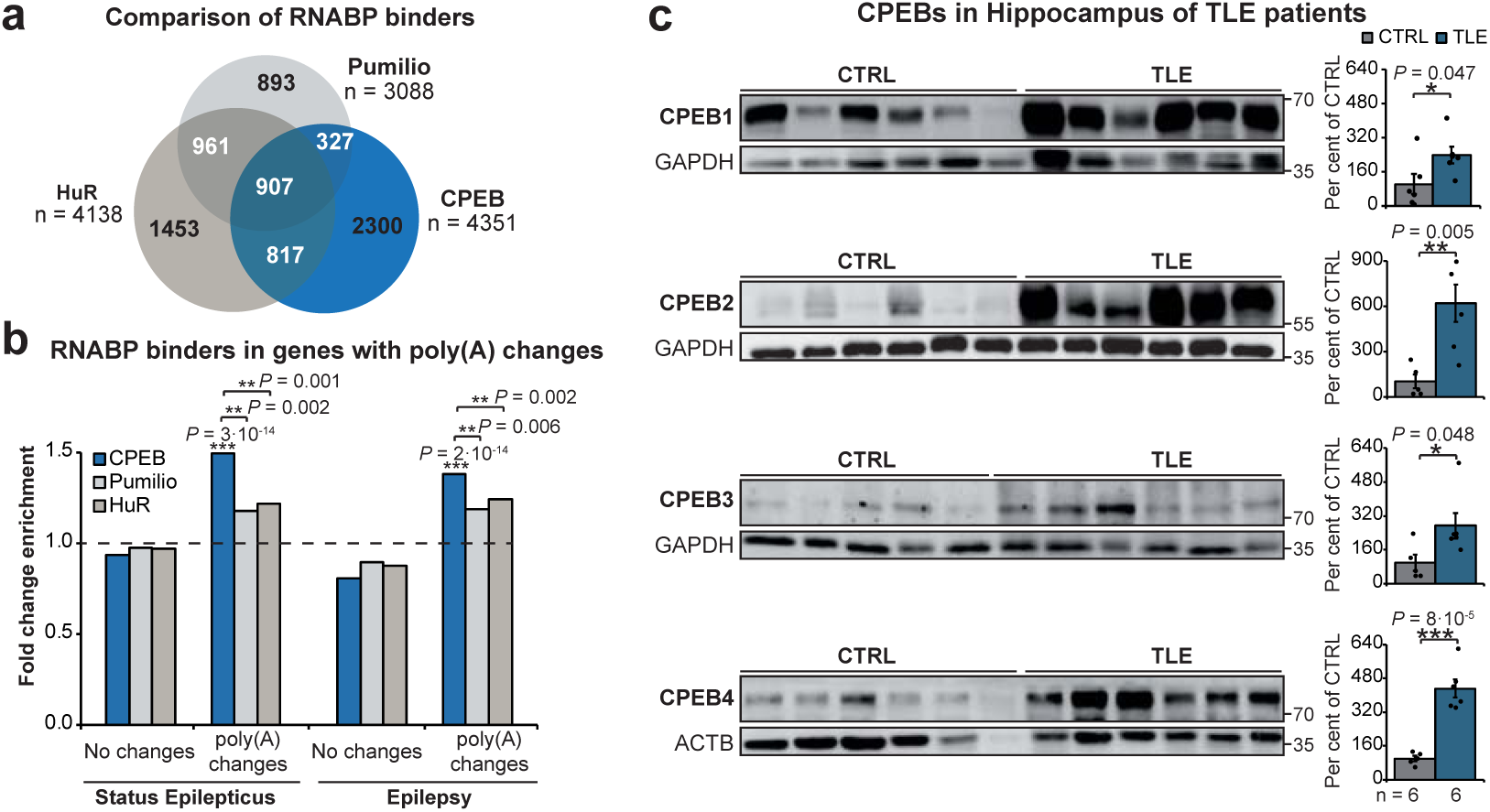
Poly(A) changes in RNABP binders and CPEBs expression in TLE. **a,** Venn diagram of identified targets of main RNABP families implicated in polyadenylation: PUMILIO, HUR and CPEBs. RNABP, RNA Binding Protein. **b,** Enrichment of RNABP binders of genes undergoing poly(A) tail changes post-status epilepticus and during epilepsy. **c,** CPEB protein levels in the hippocampus of control and patients with TLE (n = 6 per group). Protein quantity was normalized to the loading control (ACTB or GAPDH). TLE, Temporal Lobe Epilepsy; **b,** One-sided Fisher’s exact test. **c,** Two-sided unpaired t-test. Data are mean ± S.E.M. 95% CIs. \**P* < 0.05, \*\**P* < 0.01, \*\*\**P* < 0.001.

Next, to investigate the expression profile of the CPEB protein family during epilepsy, we analyzed resected hippocampal tissue obtained from drug-refractory TLE patients. Here, we found an increased expression of all CPEB family members, in particular of CPEB2 and CPEB4 (Fig. 2c).

Together, the enrichment of CPEB-binders among altered genes and the increased levels of CPEBs in TLE, indicate that members of the CPEB protein family, particularly the CPEB2-4 subfamily, are likely to be the main drivers of poly(A) tail changes during epilepsy.

### Increased expression of CPEB4 in experimental models of status epilepticus

Next, to provide further evidence of CPEB driving poly(A) tail changes during epilepsy development, we also analyzed the expression profile of the CPEB protein family in our mouse models of status epilepticus. Whereas hippocampal *Cpeb3* and *Cpeb4* transcript levels were increased at the early time points following status epilepticus in the intraamygdala KA mouse model, no changes were found in *Cpeb1* and *Cpeb2* transcription levels (Fig. 3a). At the protein level, only CPEB4 was significantly increased at short time-points, starting at 4 h post-status epilepticus and returning to baseline control levels at 24 h (Fig. 3b), the time-point at which a slight increase in CPEB1 and slight decrease in CPEB2 protein levels was also observed. Interestingly, increased transcription of *Cpeb4* was most evident in the dentate gyrus (Extended Data Fig. 3a), a subfield of the hippocampus largely resistant to seizure-induced neuronal death in the model^24^. CPEB4 protein was also upregulated in the hippocampus following status epilepticus induced by the cholinergic mimetic pilocarpine (Extended Data Fig. 3b), another widely used epilepsy model^32^. To provide additional proof of a role for CPEBs during epilepsy, we analyzed epilepsy-related genes within targets of CPEB1 and CPEB4 (Supplementary Table 2). Notably, only CPEB4 binders were enriched within epilepsy-related genes, further suggesting that CPEB4 is an important regulatory protein during epilepsy (Fig. 3c).

**Fig. 3.**
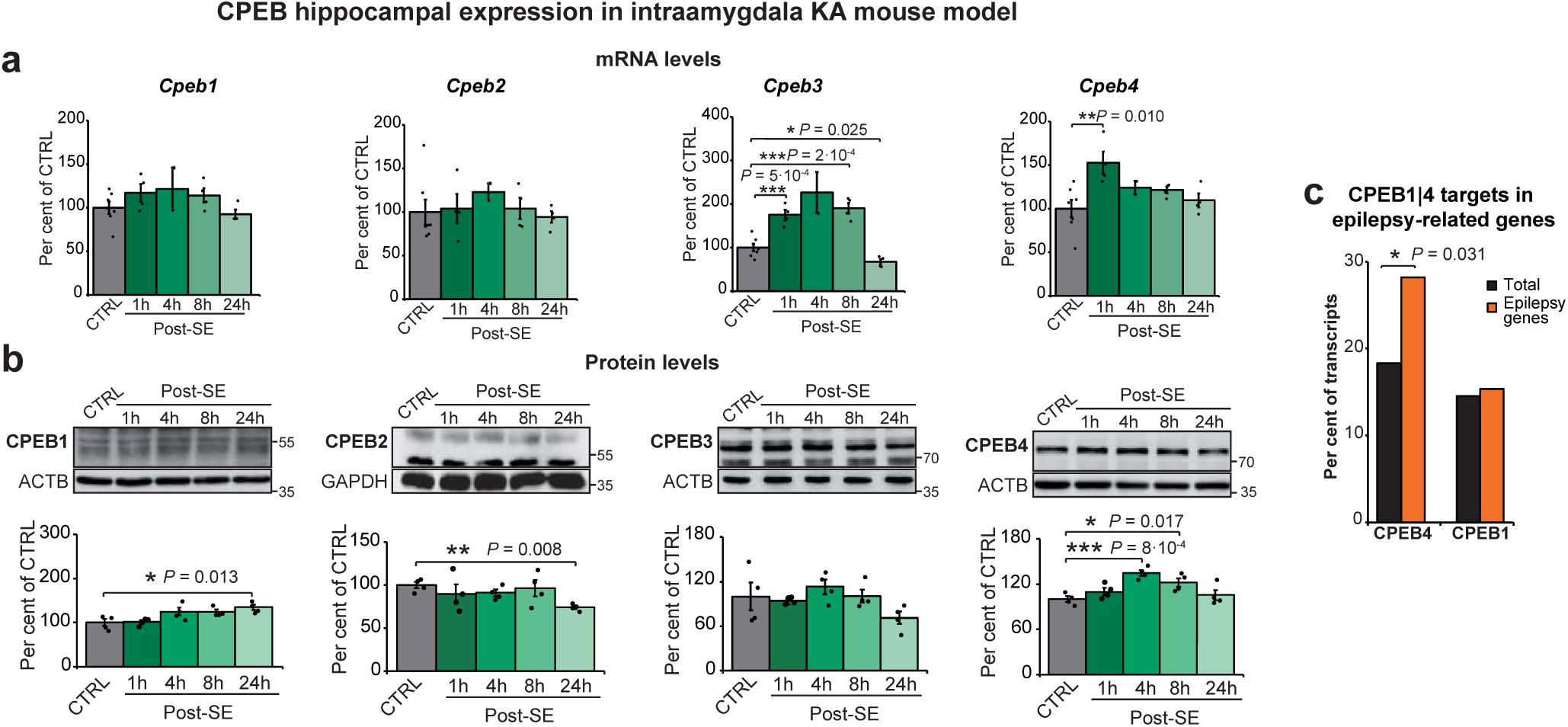
Expression of CPEBs in intraamydala KA mouse model. **a,** mRNA levels of *Cpebs* in the ipsilateral hippocampus of WT mice injected with vehicle (CTRL) vs. KA at 1 h, 4 h, 8 h and 24 h post-status epilepticus (post-SE) (n = 4). Data were analyzed and normalized to the expression of *Actb*. **b,** Protein levels of CPEB family members at same time-points (vehicle n = 7, KA n = 4). Protein quantity was normalized to the loading control (ACTB or GAPDH). **c,** Percentage of transcripts bound by CPEB4 and CPEB1 in whole transcriptome and in epilepsy-related genes. **a, b,** Two-sided unpaired t-test, **c,** One-sided Fisher’s exact test. Data are mean ± S.E.M. 95% CIs. \**P* < 0.05, \*\**P* < 0.01, \*\*\**P* < 0.001.

In summary, our data show altered expression of CPEBs in the hippocampus following status epilepticus, in particular CPEB4, with CPEB4 binders being enriched among epilepsy-related genes.

### Altered CPEB4 function may explain changes in poly(A) tail length in epilepsy

Next, to obtain functional evidence that CPEB4 is responsible for changes in mRNA polyadenylation occurring during epilepsy, we compared our identified global poly(A) profile following status epilepticus and during epilepsy to the poly(A) profile present in CPEB4 knock-out (KO) mice^21^. Notably, analysis of the global transcript polyadenylation status in CPEB4^KO/KO^ mice revealed an opposing poly(A) tail length pattern to that observed in our epilepsy mouse model (Fig. 4a). Moreover, in contrast to the observed polyadenylation signature during epilepsy, in CPEB4^KOKO^ mice the enrichment in epilepsy-related genes was found in the set of genes showing poly(A) tail lengthening (Fig. 4b). Together, these results further corroborate poly(A)-tail alterations observed in our intraamygdala KA mouse model to be attributable to altered CPEB4 function.

**Fig. 4.**
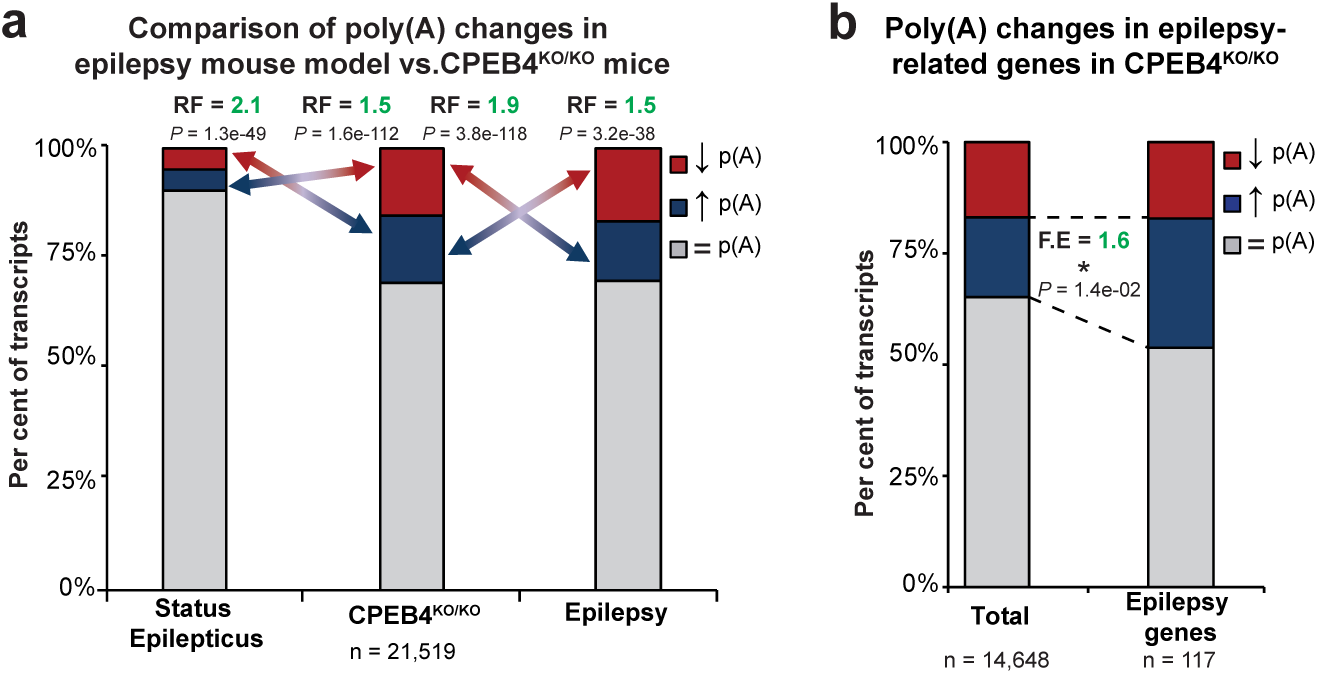
CPEB4 as main driver of poly(A) changes in epilepsy. **a,** Comparison of genes with poly(A)-tail changes in CPEB4^KO/KO^ and genes with changes in poly(A) tail length post-status epilepticus and during epilepsy; RF, representation factor; FE, fold enrichment. **b,** Percentage of epilepsy-related genes with poly(A) tail changes in CPEB4KO/KO. **a,** Hypergeometric test. **b,** One-sided Fisher’s exact test, *P* values of epilepsy-related transcripts lengthened vs. total.

### CPEB4 deficiency increases seizure susceptibility and seizure-induced brain damage

To determine whether mRNA polyadenylation and CPEB4 contribute to brain excitability or the pathophysiology of status epilepticus, we characterized seizures and their neuropathological sequelae in CPEB4 heterozygous (CPEB4^KO/+^) and homozygous (CPEB4^KO/KO^) knockout mice^33^. Immunoblot and transcript analysis confirmed a partial and full reduction of CPEB4 protein in the hippocampus of heterozygous and homozygous mice, respectively (Extended Data Fig. 4a). CPEB4-deficient mice also showed normal levels of different cell-type markers and kainate receptor levels in the hippocampus (Extended Data Fig. 4b). Furthermore, hippocampal mRNA levels of the neuronal activity-regulated gene *c-Fos* and baseline EEG recordings were similar between wildtype (WT), CPEB4^KO/+^ and CPEB4^KO/KO^ mice suggesting loss of CPEB4 does not noticeably alter normal brain function (Extended Data Fig. 4c, d).

We then investigated the impact of CPEB4-deficiency on status epilepticus triggered by an intraamygdala microinjection of KA. Both CPEB4^KO/+^ and CPEB4^KO/KO^ mice showed a shorter latency to the first seizure burst compared to WT mice (Fig. 5a). CPEB4^KO/+^ and CPEB4^KO/KO^ mice also experienced more severe seizures, as evidenced by higher total power (Fig. 5b, c) and amplitude (Extended data Fig. 4e) during the time of KA injection until the administration of the anticonvulsant lorazepam 40 min later. This increase in seizure severity persisted for an additional 60 min recording period in CPEB4^KO/KO^ mice (Fig. 5b, c). Analysis of high frequency high amplitude (HFHA) paroxysmal discharges which correlate with seizure-induced brain pathology^34^, revealed that both CPEB4^KO/+^ and CPEB4^KO/KO^ mice showed longer durations of HFHA spiking (Fig. 5d). Behavioral seizures were also more severe in CPEB4^KO/+^ and CPEB4^KO/KO^ mice during status epilepticus (Fig. 5e). Next, we analysed brain sections to determine whether loss of CPEB4 affects neuropathological outcomes. Status epilepticus in the intraamygdala model produces a characteristic lesion within the ipsilateral CA3 subfield comprising select neuron loss and gliosis. Both CPEB4^KO/+^ and CPEB4^KO/KO^ mice displayed increased neuronal death following status epilepticus as evidenced by more Fluorojade (FjB)-positive cells in all subfields of the hippocampus and in the cortex (Fig. 5f, g).

**Fig. 5.**
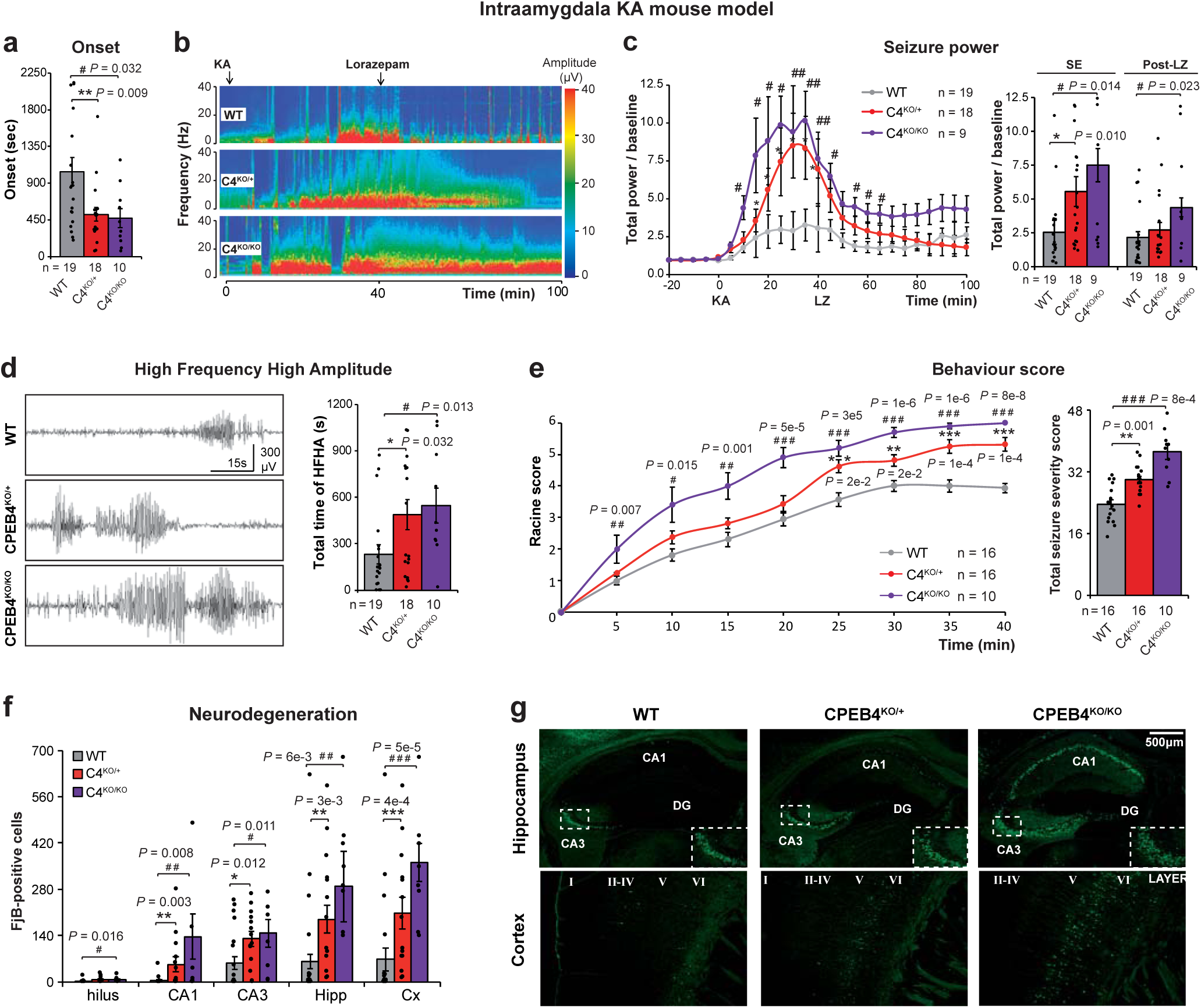
CPEB4-deficiency increases seizure susceptibility during status epilepticus and seizure-induced brain damage. **a,** Later seizure onset in CPEB4-deficient mice. **b,** Representative heatmap showing increased total seizure power during a 100 min recording period starting from intra-amygdala KA injection (t = 0). **c,** Total power normalized to baseline post-KA injection represented in 5 min segments during status epilepticus (0 – 40 min) and post-LZ administration (40 – 100 min). LZ, lorazepam. **d,** Representative electroencephalogram (EEG) traces during status epilepticus and total time of high-frequency high-amplitude (HFHA) spiking. **e,** Behavioral severity of seizures (mean Racine score) scored each 5 min and total score. **f,** Quantitative analysis of Fluorojade-B (FjB) positive cells and **g,** representative sections of neurodegeneration in ipsilateral cortex and hippocampus 72 h post-status epilepticus. Data are mean ± S.E.M. 95% CIs. WT vs CPEB4^KO/+^ \**P* < 0.05, \*\**P* < 0.01. WT vs CPEB4^KO/KO #^*P* < 0.05, ^##^*P* < 0.01, ^###^*P* < 0.001.

We also investigated neuropathological outcomes in a second model. CPEB4^KO/+^ mice subjected to status epilepticus induced by intraperitoneal pilocarpine showed increased mortality (Extended data Fig. 5a) and underwent more severe seizures (Extended data Fig. 5b, c), including longer durations of HFHA spiking (Extended data Fig. 5d). Analysis of brain sections from these mice also identified more neurodegeneration in the hippocampus and cortex of CPEB4^KO/+^ mice compared to WT controls (Extended data Fig. 5e).

Together, our results demonstrate that loss of CPEB4 increases vulnerability to seizures and hippocampal damage indicating that CPEB4 is an important regulator of brain excitability and seizure-induced neurodegeneration.

## Discussion

In the present study, we are the first to describe changes in the global mRNA polyadenylation profile following status epilepticus and during epilepsy. This represents an important post-transcriptional level of regulation of gene expression in which CPEBs play a key role. This is evidenced by bioinformatics analysis of various RNABPs and of epilepsy-related genes, together with the alteration of CPEBs in samples from human TLE and KA mouse models. We further identify CPEB4 as a key neuroprotective regulator of mRNA polyadenylation during epilepsy, and this is corroborated in CPEB4-deficient mice.

Despite the availability of over 25 anti-epileptic drugs (AEDs), pharmacological interventions remain ineffective in 30% of patients and, even when seizure-suppressive, current AEDs are merely symptomatic and have no beneficial effect on the development of epilepsy^35^. While both acute status epilepticus and chronic epilepsy are characterized by altered gene transcription, other post-transcriptional and post-translational mechanisms have been shown to be crucial during the development of epilepsy^36^. We are, however, still far from a complete picture of the pathological molecular changes occurring during epileptogenesis and epilepsy, a critical requirement for the development of much needed new therapeutic strategies. Adding to the complexity of gene regulation during epileptogenesis, we have now identified a novel regulatory gene expression mechanism, cytoplasmic mRNA polyadenylation, potentially contributing to how protein expression is controlled during epilepsy and thereby contributing to the generation of hyperexcitable networks.

Besides the long-known role of CPEB-dependent cytoplasmic mRNA polyadenylation during early development, a role has more recently been recognized in the adult brain^18^, where it is associated with synaptic plasticity, long-term potentiation, learning and memory^37^. Long forms of synaptic plasticity involving prion-like tag-mechanisms are particularly reliant on CPEB-dependent polyadenylation^38^. Epilepsy is a long lasting persistence of the hyperexcitability observed in status epilepticus and, thus, amenable to be regulated by CPEB- and poly(A)-mediated post-transcriptional regulation. In fact, while transcriptional changes are profuse during status epilepticus, they decline in epilepsy, while the percentage of genes showing altered poly(A) tail length increases from status epilepticus to epilepsy, with almost 30% of the genome affected in chronic epilepsy. These results suggest that transcriptional changes may predominate in the early phases of hyperexcitability while poly(A)-dependent regulation plays a larger role in the maintenance of epilepsy.

Despite multiple lines of evidence pointing to a key role of CPEBs in poly(A) changes in epilepsy reported here, we cannot however exclude a contribution of other RNABPs such as HUR or PUMILIO, which have previously been associated with epilepsy^39,40^. Similarly, among the various CPEBs, we show here that CPEB4 is a key player in epilepsy related poly(A)-dependent gene regulation, but we cannot discard that other CPEBs also play a role.

The strongest evidence suggesting a role for CPEB4 include its early increase in expression following acute seizures in two mouse models of status epilepticus and that a polyadenylation profile opposite to the one observed during epilepsy is observed in CPEB4 knock-out mice. Our data indicate that CPEB4 induction during epileptogenesis is neuroprotective and that the high levels of expression found in human TLE patient tissue most likely represent an endogenous antiepileptogenic adaptive mechanism to protect the brain once pathological processes are initiated. Accordingly, CPEB4 deficiency in mice, despite having no apparent impact on normal brain physiology^41^, lowers the seizure threshold upon challenge with proconvulsants and leads to an increase in seizure severity and resulting brain damage. Interestingly, further evidence of the neuroprotective role of CPEB4 was also previously obtained in an *in vitro* model of ischaemia^20^.

In conclusion, our results uncover cytoplasmic mRNA polyadenylation as an important layer of gene expression regulation in epilepsy, which must be considered when analyzing molecular patho-mechanisms during epileptogenesis and open the door for the development of novel therapies targeting mRNA polyadenylation for the treatment of drug-refractory epilepsy.

## Methods

### Human brain tissue

This study was approved by the Ethics (Medical Research) Committee of Beaumont Hospital, Dublin (05/18), and written informed consent was obtained from all patients. Briefly, patients (n = 6) were referred for surgical resection of the temporal lobe for the treatment of intractable TLE. After temporal lobe resection, hippocampi were obtained and frozen in liquid nitrogen and stored at −70°C until use. A pathologist (Dr. Michael Farrell) assessed hippocampal tissue and confirmed the absence of significant neuronal loss. Control (autopsy) temporal hippocampus (n = 6) was obtained from individuals from the Brain and Tissue Bank for Developmental Disorders at the University of Maryland, Baltimore, MD, U.S.A. Samples were processed for Western blot analysis. Brain sample and donor metadata are available in Supplementary Table 4.

### Animal models of status epilepticus and epilepsy

Animal experiments were carried out in accordance with the principles of the European Communities Council Directive (2010/63/EU). Procedures were reviewed and approved by the Research Ethics Committee of the Royal College of Surgeons in Ireland (REC 1322) and the Irish Health Products Regulatory Authority (HPRA) (AE19127/P038). All efforts were maximized to reduce the number of animals used in this study. Mice used in our experiments were 8-12 weeks old male and female C57Bl/6, obtained from Harlan Laboratories (Bicester, UK) and from the Biomedical Research Facility (BRF), Royal College of Surgeons in Ireland (Dublin, Ireland). CPEB4-deficient mice (CPEB4^KO/+^, CPEB4^KO/KO^) harbour a heterozygous or homozygous deletion of constitutive exon 2, respectively, resulting in a premature stop codon and thereby a partial or full suppression of CPEB4 protein expression^33^. Animals were housed in a controlled biomedical facility on a 12 h light/dark cycle at 22±1 °C and humidity of 40-60% with food and water provided *ad libitum*. Status epilepticus was induced as described previously either via an intraamygdala injection of KA or an intraperitoneal injection of pilocarpine^42^. Before implantation of cannulas (intraamygdala KA injection) and electrodes (EEG recordings), mice were anesthetized using isoflurane (5% induction, 1-2% maintenance) and maintained normothermic by means of a feedback-controlled heat blanket (Harved Apparatus Ltd, Kent, UK). Next, mice were placed in a stereotaxic frame and a midline scalp incision was performed to expose the skull. A guide cannula (coordinates from Bregma; AP = −0.94 mm, L = −2.85 mm) and three electrodes (Bilaney Consultants Ltd, Sevenoaks, UK), two above each hippocampus and one above the frontal cortex as reference, were fixed in place with dental cement. Intraamygdala KA (0.3 µg KA in 0.2 µl phosphate-buffered saline (PBS)) (Sigma-Aldrich, Dublin, Ireland) was administered into the basolateral amygdala nucleus. Vehicle-injected control animals received 0.2 µl of PBS. To reduce morbidity and mortality, mice were treated with an i.p. injection of the anticonvulsant lorazepam (6 mg/kg) 40 min post-KA or PBS injection (Wyetch, Taplow, UK). As described previously, all mice develop epilepsy after a short latency period of 3-5 days^24^. Status epilepticus was also induced by an i.p. injection of pilocarpine (340 mg/kg body weight) (Sigma-Aldrich, Dublin, Ireland) 20 min following the injection of methyl-scopolamine (1 mg/kg) (Sigma-Aldrich, Dublin, Ireland)^42^. Mice were treated with i.p. lorazepam (6 mg/kg) 90 min following i.p. pilocarpine. EEG was recorded using the Xltek recording system (Optima Medical Ltd, Guildford, UK) starting 20 min before administration of pro-convulsant (KA or pilocarpine) to record baseline and during status epilepticus (40 min for intraamygdala KA-treated mice and 90 min for mice treated with i.p. pilocarpine). EEG was continued for 1 h post-administration of lorazepam. Mice were euthanized at different time-points following status epilepticus (1 h, 4 h, 8 h, 24 h) and during chronic epilepsy (14 days) and brains flash-frozen whole in 2-methylbutane at −30 °C for FjB staining, perfused with PBS and paraformaldehyde (PFA) 4% for immunofluorescence or microdissected and frozen for protein or RNA extraction.

### Poly(U) chromatography

C57Bl/6 WT mice were sacrificed 8 h following status epilepticus (acute pathology) (KA and PBS, n = 9 per group) or 14 days post-status epilepticus (time-point at which all mice suffer from chronic epilepsy^24^) (KA and PBS, n = 9 per group). Ipsilateral hippocampi were quickly dissected, pooled into 3 groups (n = 3 per pooled sample) and stored at −80°C until use. RNA was extracted using the Maxwell® 16 LEV simplyrna Tissue Kit (Promega, AS1280). Poly(A) RNA fraction was purified by poly(U) chromatography as before^43^. Poly(U)-agarose (Sigma, p8563) was suspended in swelling buffer (0.05 M Tris-HCl, pH 7.5, 1 M NaCl) 35 ml/g, incubated overnight at room temperature and loaded into a chromatography column. An aliquot of total RNA was stored at −80°C (“Input”) and the remaining sample incubated with sample buffer (0.01 M Tris-HCl, pH 7.5, 1 mM EDTA, 1% SDS) for 5 min at 65°C and chilled on ice. Binding buffer was added (0.05 M Tris-HCl, pH 7.5, 0.7 M NaCl, 10 mM EDTA, 25% [v/v] formamide) which was followed by loading of samples into the poly(U)-agarose chromatography column (Mobitec, M1002s). Samples were then incubated for 30 min at room temperature (25°C) with agitation. Next, columns containing samples were washed three times at 25°C and six times at 55°C with washing buffer (0.05 M Tris-HCl, pH 7.5, 0.1 M NaCl, 10 mM EDTA, 25% [v/v] formamide). The 55°C washes were collected and stored at −80°C (“Short poly(A)-tail fraction”). The remaining poly(A) RNA (“Long poly(A)-tail fraction”) was eluted with elution buffer (0.05 M HEPES, pH 7, 10 mM EDTA, 90% [v/v] formamide) at 55°C and stored at −80°C. RNA of the two poly(A) fractions was precipitated by adding 1 volume of isopropanol, 1/10^th^ volume of sodium acetate 3 M pH 5.2 and 20 µg of glycogen (Sigma, G1767). Samples were incubated at −20°C for 20 min and centrifuged for 15 min at 14000 g at 4°C. Supernatant was removed and pellet was washed with 750 µl of ethanol and centrifuged at 14000 g at 4°C for 5 min. Then supernatant was removed again and pellet was air-dried for 5 min. Next, RNAs were resuspended in 300 µl of nuclease-free water and 300 µl of acid Phenol:Chloroform (5:1) were added. Then, samples were vortexed and centrifuged for 10 min at 14000 g at 4°C. The aqueous phase was recovered, mixed with 1 volume of chloroform, vortexed and centrifuged again. The aqueous phase was recovered and precipitated again using the isopropanol precipitation. To confirm the average length in each fragment, when setting up the method, we performed digestion of the non-poly(A) mRNA regions followed by end-labelling of the poly(A) tail for each eluted fraction and Urea-PAGE. Ensuring proper functioning of our technique, poly(A)-tail changes were assessed by HIRE-PAT assays of control genes in Input, Washed and Eluted fractions. RNA quantification was performed by Qubit Fluorimeter using Qubit RNA Hs Assay kit (Thermo-Fisher Scientific, Q32852). RNA integrity QC was performed with Agilent Bioanalyzer 2100, using RNA Nano Assay (Agilent Technologies 5067-1511) and RNA Pico Assay (Agilent Technologies 5067-1513).

### GeneAtlas MG-430 PM microarray analysis

cDNA library preparation and amplification were performed according to the manufacturer’s instructions (Sigma-Aldrich) using the WTA2 kit with 25 ng starting material. cDNA was amplified for 17 cycles and purified using PureLink Quick PCR Purification Kit (Invitrogen, K310001). Quantification of amplified cDNA was carried out on a Nanodrop ND-1000 spectrophotometer (Thermo-Fisher Scientific, Waltham, MA, USA). 8.5 µg of the cDNA from each sample were fragmented and labelled with GeneChip Mapping 250K Nsp assay kit (Affymetrix, 900753) following the manufacturer’s instructions. Hybridization was performed using the GeneAtlas Hyb, Wash and Stain Kit for 3’ IVT arrays. Samples ready to hybridize were denatured at 96°C for 10 min prior to incubation with mouse MG-430 PM Array Strip (Affymetrix, 901570). Hybridization was performed for 16 h at 45°C in the GeneAtlas Hybridization Oven (Affymetrix, 00-0331). Washing and staining steps after hybridization were performed in the GeneAtlas Fluidics Station (Affymetrix, 00-0079), following the specific script for Mouse MG-430 PM Arrays. Finally, arrays were scanned with GeneAtlas Scanner (Affymetrix) using default parameters, and the generation of CEL files for bioinformatics analysis was performed using GeneAtlas software (Affymetrix). Processing of microarray samples was performed using R^44^ and Bioconductor^45^. Raw CEL files were normalized using RMA background correction and summarization^46^. Standard quality controls were performed in order to identify abnormal samples^47^ regarding: a) spatial artefacts in the hybridization process (scan images and pseudo-images from probe level models); b) intensity dependences of differences between chips (MvA plots); c) RNA quality (RNA digest plot); and d) global intensity levels (boxplot of perfect match log-intensity distributions before and after normalization and RLE plots). Probeset annotation was performed using the information available on the Affymetrix web page (https://www.affymetrix.com/analysis/index.affx) using version na35. Expression values were adjusted for technical biases as described^48^ using a linear model and implemented with the R package “limma”^49^. For each biological replicate the log2 fold change was computed between “WASHED (Short)” and “ELUTED (Long)” samples and used to find significant differences between intraamygdala KA or vehicle-treated control mice sacrificed at two different time-points (8 h (acute) and 14 days (chronic epilepsy) following status epilepticus). Differential expression was performed using a linear model with fluidics and amplification batch as covariates. *P*-values were adjusted with the Benjamini and Hochberg correction. We considered one transcript as shortened when *P*-value was < 0.05 and FC was negative and lengthened when *P*-value was < 0.05 and FC was positive, in at least one probe. If the same transcript showed opposite results for different probes, the poly(A) tail was considered as not changed.

### Gene Ontology analysis

Genes with changes in poly(A) tail length between intraamygdala KA or vehicle control mice sacrificed at two different time-points (8 h and 14 days following status epilepticus) were analysed by GO terms with the bioinformatic tool DAVID Bioinformatics Resources 6.7^25^.

### High-Resolution poly(A) tail (HIRE-PAT) assay

To measure poly(A) tail length of mRNAs, USB® Poly(A) Tail-Length Assay Kit (Affymetrix, 76455) based on the HIRE-PAT method was performed. We used total RNA obtained of Washed and Eluted fractions (enriched in mRNA short and long poly(A) tail respectively), from the ipsilateral hippocampi of mice sacrificed 8 h following status epilepticus. G/I tailing (1 μg of total RNA) and reverse transcription were performed according to the manufacturer’s instructions. Poly(A) tail size was determined by subtracting the PCR amplicon size obtained with the Universal primer and forward specific primers. To verify that the measured poly(A) tail corresponds to a specific gene, at least two different forward specific primers were tested (Supplementary Table 5). PCR products were resolved on a 2.5% sybr green agarose gel (Biotium, 41004) run at 120 V for 1.5 h.

### Western blot

Western blot was carried out as before^42^. Samples (mouse and human) were prepared by homogenizing extracted brain tissue in ice-cold extraction buffer (20 mM HEPES pH 7.4, 100 mM NaCl, 20 mM NaF, 1% Triton X-100, 1 mM sodium orthovanadate, 1 μM okadaic acid, 5 mM sodium pyrophosphate, 30 mM β-glycerophosphate, 5 mM EDTA, protease inhibitors (Complete, Roche, Cat. No 11697498001)). Protein concentration was determined by Quick Start Bradford kit assay (Bio-Rad, 500-0203) or BCA (Thermo Scientific, Ref 23235) following the manufacturer’s instructions. Between 15 - 40 μg of total protein were electrophoresed on 8 - 10% sodium dodecyl sulfate (SDS)-polyacrylamide gel, transferred to a nitrocellulose blotting membrane (Amersham Protran 0.45 μm, GE Healthcare Life Sciences, 10600002) and blocked in TBS-T (150 mM NaCl, 20 mM Tris–HCl, pH 7.5, 0.1% Tween 20) supplemented with 5% non-fat dry milk. Membranes were incubated overnight at 4°C with the primary antibody in TBS-T supplemented with 5% non-fat dry milk. On the next day, following washing with TBS-T, membranes were incubated with secondary HRP-conjugated anti-mouse or anti-rabbit IgG (1:1000, Jackson Immuno 266 Research, Plymouth, PA, U.S.A), or anti-goat IgG-Fc fragment (1:5000, Bethyl, A50-104P) and protein bands visualized using chemiluminescence Merck Millipore, Billerica, MA, U.S.A (Pierce Biotechnology, Rockford, IL, USA). Gel bands were captured using a Fujifilm LAS-4000 (Fujifilm, Tokyo, Japan), analysed using Alpha-EaseFC4.0 software and quantified using ImageJ software^50^. Protein quantity was normalized to the loading control (ACTB or GAPDH). The following primary antibodies were used: rabbit GRIN2B (1:1000, Abcam, ab65783); rabbit METTL3 (1:1000, Abcam, ab195352); mouse STX6 (1:1000, BD transduction Laboratories, Cat 610635); rabbit CPEB1 (1:350, Santacruz, sc-33193, for human samples and 1:1000, proteintech, 13274-1-AP for mouse samples); rabbit CPEB2 (1:1000, Abcam, ab51069); rabbit CPEB3 (1:1000, Abcam, ab10883); rabbit CPEB4 (1:1000, Abcam, ab83009); rabbit NEUN (1:1000, Millipore, ABN78); mouse GFAP (1:1000, Sigma, G3893); rabbit GLUR6/7 (1:1000, Millipore, 1497226); mouse ACTB (1:5000, Sigma, A2228); mouse GAPDH (1:5000, Cell Signaling Technology, 14C10);

### RNA extraction and quantitative polymerase chain reaction (qPCR)

RNA extraction was performed using the Trizol method, as described before^42^. Quantity and quality of RNA was measured using a Nanodrop Spectrophotometer (Thermo Scientific, Rockford, IL, U.S.A). Samples with a 260/280 ratio between 1.8 – 2.2 were considered acceptable. 500 ng of total mRNA was used to produce complementary DNA (cDNA) by reverse transcription using SuperScript III reverse transcriptase enzyme (Invitrogen, CA, U.S.A) primed with 50 pmol of random hexamers (Sigma, Dublin, Ireland). qPCR was performed using the QuantiTech SYBR Green kit (Qiagen Ltd, Hilden, Germany) and the LightCycler 1.5 (Roche Diagnostics, GmbH, Mannheim, Germany). Each reaction tube contained 2 μl cDNA sample, 10 μl SyBR green Quantitect Reagent (Quiagen Ltd, Hilden, Germany), 1.25 μM primer pair (Sigma, Dublin, Ireland) and RNAse free water (Invitrogen, CA, U.S.A) to a final volume of 20 μl. Using LightCycler 1.5 software, data were analysed and normalized to the expression of *Actb*. Primers used are detailed in Supplementary Table 5.

### Enrichment analysis

To evaluate whether a gene set is enriched over the background, enrichment analysis studies were carried out using one-sided Fisher’s exact test. For our analysis, we used curated gene lists of epilepsy-related genes generated from three independent studies. This included genes with mutation that cause epilepsy (n = 84); mutations in these genes cause pure or relatively pure epilepsies, or syndromes with epilepsy as the core symptom^26^. Genes with ultra-rare deleterious variation in familial genetic generalized epilepsy (GGE) and non-acquired focal epilepsy (NAFE) (n = 18); we chose the most significant genes per group (top 15)^27^. Genes localized in loci associated with epilepsy (n = 21); the 21 most likely epilepsy genes at these loci, with the majority in genetic generalized epilepsies, were selected^28^. The complete set of epilepsy-related genes used in our study is shown in Supplementary Table 2.

### Identification of mRNA targets of RNA binding proteins

CPEB1 and CPEB4 binders were determined previously by RNA immunoprecipitation^21^; PUM1 and PUM2 targets were identified by individual nucleotide resolution CLIP (iCLIP)^31^; genes that interact with HUR were identified by photoactivatable ribonucleoside-enhanced crosslinking and immunoprecipitation (PAR-CLIP)^30^. The complete list of RNABP targets is shown in Supplementary Table 3. The list of brain-specific genes was obtained from the human protein atlas (http://proteinatlas.org/humanproteome/brain).

### Electroencephalogram (EEG) analysis

To analyse EEG frequency, amplitude signal (power spectral density and EEG spectrogram of the data) and seizures onset, EEG data was uploaded into Labchart7 software (AD instruments Ltd, Oxford, UK). EEG total power (μV^2^) is a function of EEG amplitude over time and was analysed by integrating frequency bands from 0 to 100 Hz. Power spectral density heat maps were generated using LabChart (spectral view), with the frequency domain filtered from 0 to 40 Hz and the amplitude domain filtered from 0 to 50 mV. The duration of high-frequency (>5 Hz) and high-amplitude (>2 times baseline) (HFHA) polyspike discharges of ≥5 s duration, synonymous with injury-causing electrographic activity^34^, were counted manually by a reviewer who was blinded to treatment. Seizure onset was calculated as first seizure burst from time of intraamygdala KA injection.

### Behaviour assessment of seizure severity

Changes in seizure-induced behaviour were scored according to a modified Racine Scale as reported previously^51^. Score 1, immobility and freezing; Score 2, forelimb and or tail extension, rigid posture; Score 3, repetitive movements, head bobbing; Score 4, rearing and falling; Score 5, continuous rearing and falling; Score 6, severe tonic–clonic seizures. Mice were scored by an observer blinded to treatment every 5 min for 40 min after KA injection. The highest score attained during each 5 min period was recorded.

### Fluoro-Jade B staining

To assess status epilepticus-induced neurodegeneration, FjB staining was carried out as before^52^. Briefly, 12 μm coronal sections at the medial level of the hippocampus (Bregma AP = −1.94 mm) were cut on a cryostat. Tissue was fixed in formalin, rehydrated in ethanol, and then transferred to a 0.006% potassium permanganate solution followed by incubation with 0.001% FjB (Chemicon Europe Ltd, Chandlers Ford, UK). Sections were mounted in DPX mounting solution. Then, using an epifluorescence microscope, cells including all hippocampal subfields (dentate gyrus (DG), CA1 and CA3 regions) and cortex were counted by a person unaware of treatment under a 40x lens in two adjacent sections and the average determined for each animal.

### Data analysis

Statistical analysis was performed using SPSS 21.0 (SPSS® Statistic IBM®). Data are represented as mean ± S.E.M. (Standard Error of the Mean) with 95% confidence interval. The normality of the data was analysed by Shapiro-Wilk test (n < 50) or Kolmogorov-Smirnov (n > 50). Homogeneity of variance was analysed by Levente test. For comparison of two independent groups, two-tail unpaired t-Student’s test (data with normal distribution), Mann-Whitney-Wilcoxon or Kolmogorov-Smirnov tests (with non-normal distribution) was performed. To compare dependent measurements, we used a paired t-test (normal distribution) or Wilcoxon signed-rank tests (non-normal). For multiple comparisons, data with a normal distribution were analysed by one way-ANOVA test followed by a Tukey’s or a Games-Howell’s post-hoc test. Statistical significance of non-parametric data for multiple comparisons was determined by Kruskal-Wallis one-way ANOVA test. Enrichment tests were carried out by using one-sided Fisher’s exact test. A critical value for significance of *P* < 0.05 was used throughout the study.

## Data availability

The data that support the findings of this study are available from the corresponding authors upon reasonable request. All records have been approved and assigned GEO accession numbers, however, until the acceptance of the manuscript, these will be kept private. GSE132523 - Identification of the mRNA polyadenylation profile in the hippocampus following intraamygdala kainic acid-induced of status epilepticus in mice

## Acknowledgments

This work was supported by funding from the Health Research Board HRA-POR-2015-1243 to TE; Science Foundation Ireland (13/SIRG/2098 and 17/CDA/4708 to T.E and co-funded under the European Regional Development Fund and by FutureNeuro industry partners 16/RC/3948 to D.C.H); from the H2020 Marie Skłodowska-Curie Actions Individual Fellowship (796600 to L.D-G and 753527 to E.B), from the European Union’s Horizon 2020 research and innovation programme under the Marie Sklodowska-Curie grant agreement (No. 766124 to T.E.), from ISCIII-CiberNed-PI2013/09 and SAF2015-65371-R and RTI2018-096322-B-I00 to J.J.L., A.P. was recipient of a CIBERNED-Ayuda a la movilidad. We thank the following core facilities: IRB-Functional Genomic and IRB-Bioinformatics/Biostatistics.

## Author contributions

A.P. performed bioinformatics and statistical analysis, L.D.-G. performed and was involved in most of experiments, and A.P. and L.D.-G contributed to study design and were involved in all assays and data collection. M.A. performed Western-Blot, qRT-PCR and EEG and data analysis. E.B. performed mouse modelling and behavioural studies. G.C. contributed to the microarray analysis. J.M. analysed data. I.O. performed semiquantitative qRT-PCR. Y.H.-S carried out Western-Blot and qRT-PCR analysis. N.D., M.A.F. and D.F.O. provided human tissue. D.C.H. and R.M. made intellectual contributions to experimental design and discussion. J.J.L. and T.E. directed and conceived the study. T.E. wrote the paper with input from all authors.

