## Extended Data Figures for "Large-scale changes to mRNA polyadenylation in temporal lobe epilepsy"

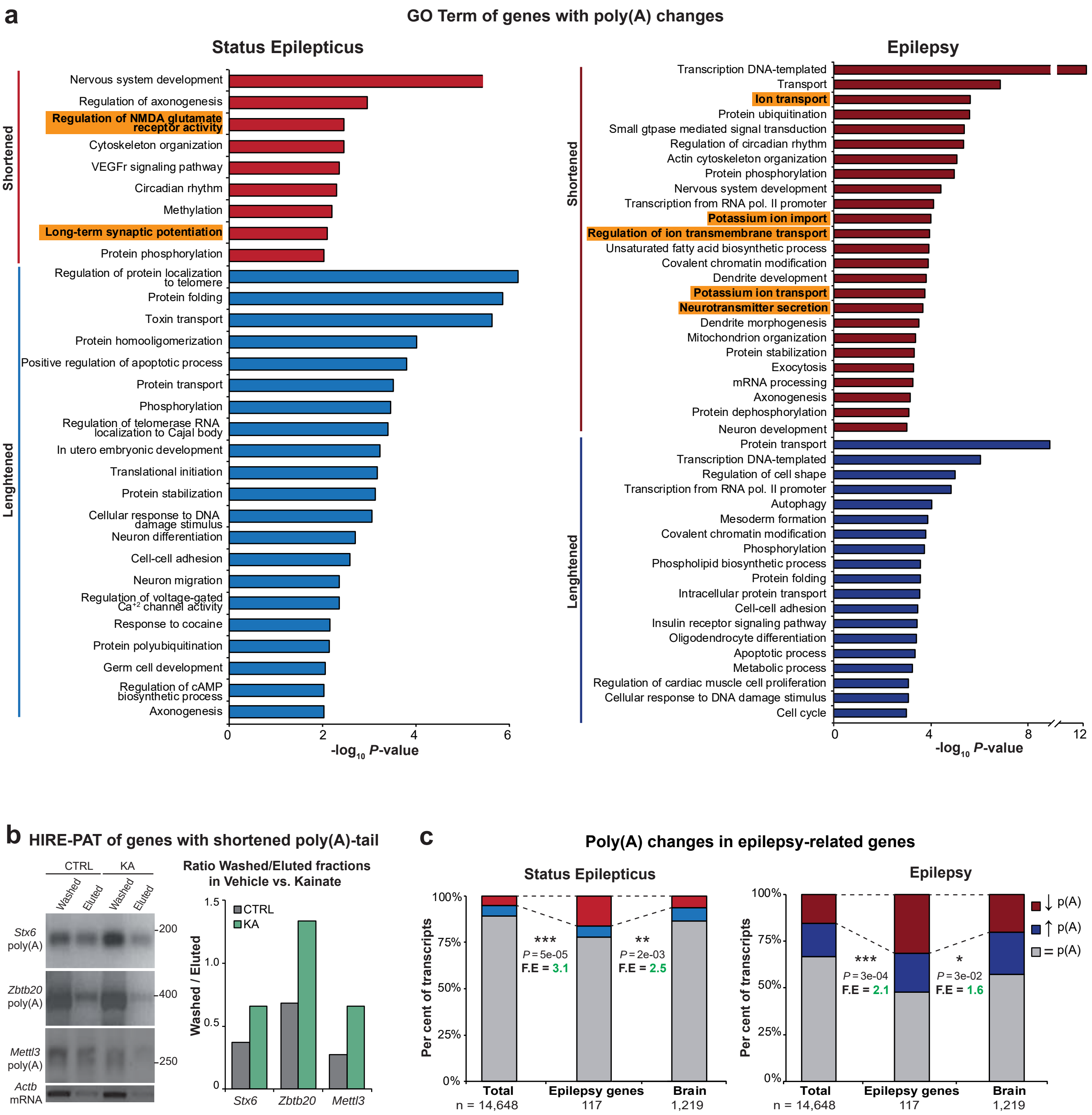


**Extended Data Fig. 1 | Supplementary information on mRNA polyadenylation and deadenylation of epilepsy-related genes**
**a,** Top GO terms associated with biological process using DAVID resources of transcripts with poly(A) tail changes following status epilepticus and during epilepsy; the negative log10 of the *P* value is plotted on the X-axis. GO, Gene Ontology **b,** HIRE-PAT assay of selected deadenylated transcripts in washed and eluted hippocampal fractions of WT mice treated with intraamygdala vehicle (CRTL) vs. KA (8 h post-status epilepticus) (n = 9 per group). **c,** Percentage of genes presenting changes in poly(A) tail length following intraamygdala KA-induced status epilepticus and during epilepsy in whole and brain-specific transcriptomes, and in epilepsy-related genes. **c,** One-sided Fisher's exact test, *P* values of deadenylated epilepsy-realted genes vs. whole and brain transcriptome. *$P < 0.05$, **$P < 0.01$, ***$P < 0.001$.

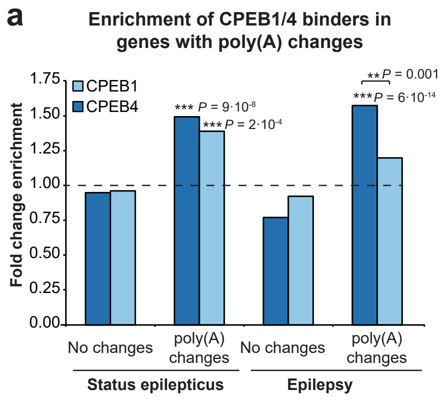


**Extended Data Fig. 2 | Poly(A) changes in CPEB1/4 binders**

**a,** Enrichment analysis of CPEB1- and CPEB4-only binders in transcripts with poly(A) changes post-status epilepticus and during epilepsy. One-sided Fisher's exact test. $^{**}P < 0.01$, $^{***}P < 0.001$.

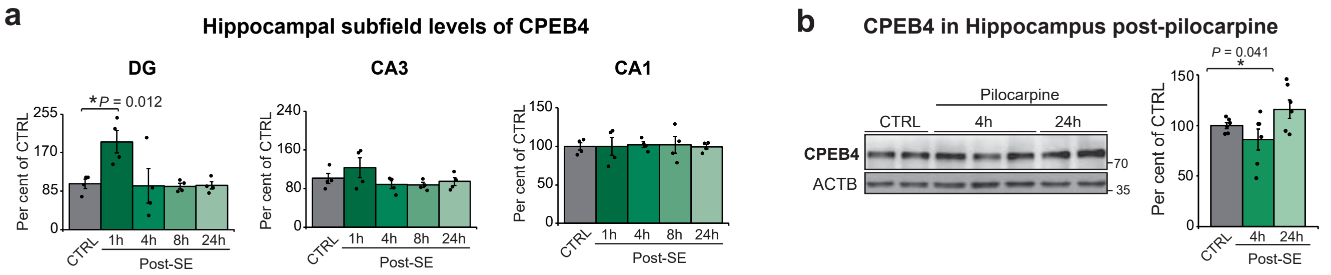


**Extended Data Fig. 3 | Hippocampal CPEB4 expression post-status epilepticus.**
**a,** mRNA levels of *Cpeb4* in the ipsilateral hippocampal subfields CA1, CA3 and DG at different time points post-intraamygdala KA-induced status epilepticus (n = 4). DG, dentate gyrus. Data were analyzed and normalized to the expression of *Actb*. **b,** CPEB4 protein levels in the hippocampus of WT mice 4 h and 24 h following intraperitoneal pilocarpine-induced status epilepticus (vehicle n = 9, pilocarpine n = 6). Protein quantity was normalized to the loading control (ACTB). **a, b,** Two-sided unpaired t-test. Data are mean ± S.E.M. 95% CIs. *$P < 0.05$

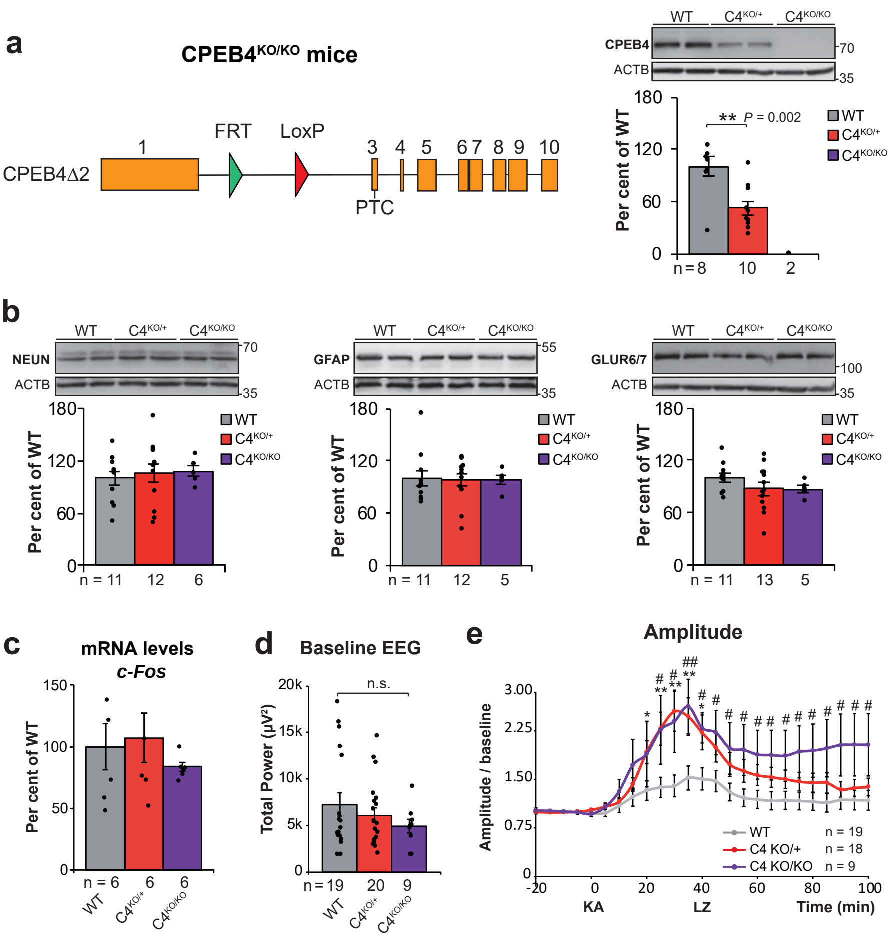


**Extended Data Fig. 4 | Supplementary information of CPEB4-deficient mice.**

**a,** Construct design and CPEB4 protein levels in the hippocampus of CPEB4$^{KO/+}$ and CPEB4$^{KO/KO}$ mice. Protein quantity was normalized to the loading control (ACTB) **b,** Protein levels of NEUN, GFAP and GLUR6/7. Protein quantity was normalized to the loading control (ACTB) **c,** *c-Fos* mRNA levels in the hippocampus of WT, CPEB4$^{KO/+}$, CPEB4$^{KO/KO}$ mice. Data were analyzed and normalized to the expression of *Actb*. **d,** Baseline EEG recordings of different genotypes. EEG, electroencephalogram **e,** Amplitude post-intraamygdala KA injection normalized to baseline. KA, kainic acid; LZ, lorazepam. Data are mean ± S.E.M. 95% CIs. WT vs CPEB4$^{KO/+}$ **P* < 0.05, ***P* < 0.01. WT vs CPEB4$^{KO/KO}$ $^{\#}P$ < 0.05, $^{\#\#}P$ < 0.01--

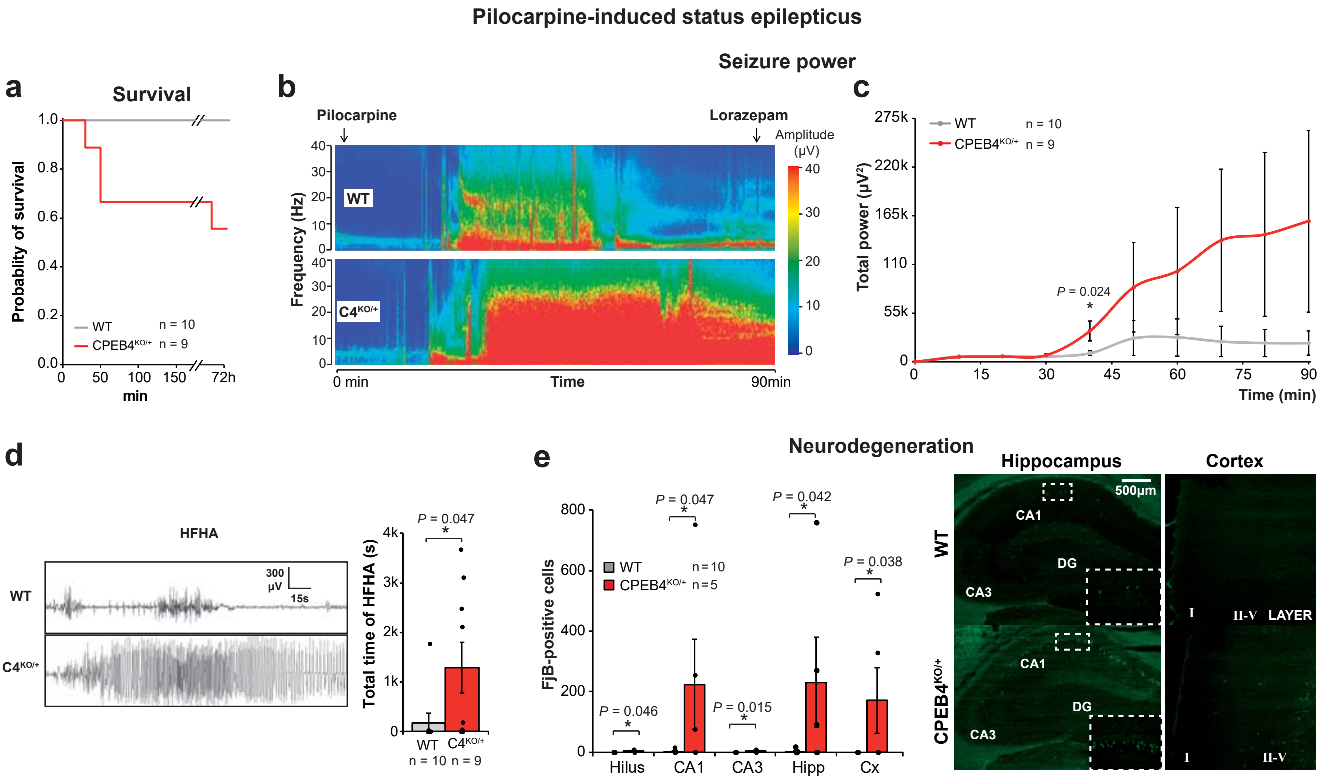


**Extended Data Fig. 5 | CPEB4-deficiency aggravates pilocarpine-induced seizures and neurodegeneration.**
**a,** Kaplan–Meier curve for cumulative survival. **b,** Representative heatmaps showing increased total seizure power in CPEB4$^{KO/+}$ during a 90 min recording period starting at intraperitoneal pilocarpine injection. **c,** Total power measured in 10 min segments. **d,** Total time of high-frequency high-amplitude (HFHA) polyspike discharges and representative EEG traces. **e,** Quantitative analysis of FluoroJade-B (FjB) -positive cells and representative sections of neurodegeneration in hippocampus and cortex 72h post-status epilepticus. Data are mean ± S.E.M. 95% CIs. *$P < 0.05$
